# GeneWalk identifies relevant gene functions for a biological context using network representation learning

**DOI:** 10.1101/755579

**Authors:** Robert Ietswaart, Benjamin M. Gyori, John A. Bachman, Peter K. Sorger, L. Stirling Churchman

## Abstract

The primary bottleneck in high-throughput genomics experiments is identifying the most important genes and their relevant functions from a list of gene hits. Existing methods such as Gene Ontology (GO) enrichment analysis provide insight at the gene set level. For individual genes, GO annotations are static and biological context can only be added by manual literature searches. Here, we introduce GeneWalk (github.com/churchmanlab/genewalk), a method that identifies individual genes and their relevant functions under a particular experimental condition. After automatic assembly of an experiment-specific gene regulatory network, GeneWalk quantifies the similarity between vector representations of each gene and its GO annotations through representation learning, yielding annotation significance scores that reflect their functional relevance for the experimental context. We demonstrate the use of GeneWalk analysis of RNA-seq and nascent transcriptome (NET-seq) data from human cells and mouse brains, validating the methodology. By performing gene- and condition-specific functional analysis that converts a list of genes into data-driven hypotheses, GeneWalk accelerates the interpretation of high-throughput genetics experiments.

## Introduction

High-throughput functional genomics experiments generate genome-scale datasets that require computational analyses^1–5^, which yield lists of ‘hit’ genes^2^. Such lists typically include hundreds to thousands of genes of interest, ranked by p-values, whose biological significance (as opposed to technical validity) is not always clear^6^. The main bottleneck is in determining which genes, and which specific functions of those genes, are most relevant to the biological context, thus prioritizing those genes worthy of further study. GO annotations are commonly used to add information related to biological processes, molecular functions and cellular components to genes and gene lists^7^, but they do not account for variations in gene function across biological contexts. This limitation is particularly notable for genes that are involved in many different processes, e.g. *EGFR*, which affects transcription, signal transduction, cell division, survival, motility, and other processes^7^. Alternatively, GO and gene set enrichment analysis (GSEA)^1,3–5,8,9^ can be used to reveal which biological processes are enriched under each specific condition, but do not address the context-specific functions of individual genes. Accordingly, new methods are required to generate functional information about individual genes under particular conditions of interest or biological contexts. To address this need, we developed GeneWalk, a knowledge-based machine learning and statistical modelling method that highlights the gene functions that are relevant for a specific biological context.

GeneWalk takes advantage of two recent advances in computational biology: deep learning to condense information^10–13^, and generation of biological networks derived from database aggregation efforts^9,14–17^. Unsupervised representation learning by neural networks can reduce the dimensionality of complex datasets or networks (graphs)^10,11^. Thus, vertices (nodes) in any network can be represented by vectors of low dimensionality informed through the network topology^12,18–20^. Networks of biological mechanisms are now available from knowledge bases^9,17^, such as Pathway Commons^14^, String^9^, Omnipath^17^, and the Integrated Network and Dynamical Reasoning Assembler (INDRA^15,16^). INDRA reaction statements (e.g., protein phosphorylation, transcriptional regulation, or biological process regulation) are extracted from the body of biomedical literature using either natural language processing systems of primary literature in the minable NCBI corpus or queries on pathway databases.

GeneWalk automatically assembles a biological network from a knowledge base and GO ontology starting with a list of genes of interest (e.g. differentially expressed genes or hits from a genetic screen) as input (Figure 1A). The network structure is learned through random walks using an unsupervised network representation learning algorithm (DeepWalk^10^). The resultant vector representations enable a quantitative comparison between genes and GO terms, highlighting the GO terms most relevant for the biological context under study. We demonstrate the applicability of GeneWalk by using it to analyze three experiments in which the data were obtained by either RNA-seq or native elongating transcript sequencing (NET-seq), which probes the nascent transcriptome. GeneWalk identified context-relevant GO terms while filtering out the majority of irrelevant GO terms for each gene, allowing the researcher to quickly hone in on relevant targets. Thus, GeneWalk serves as a rapid data-driven hypothesis-generating tool for exploratory functional gene analysis.

**Figure 1.**
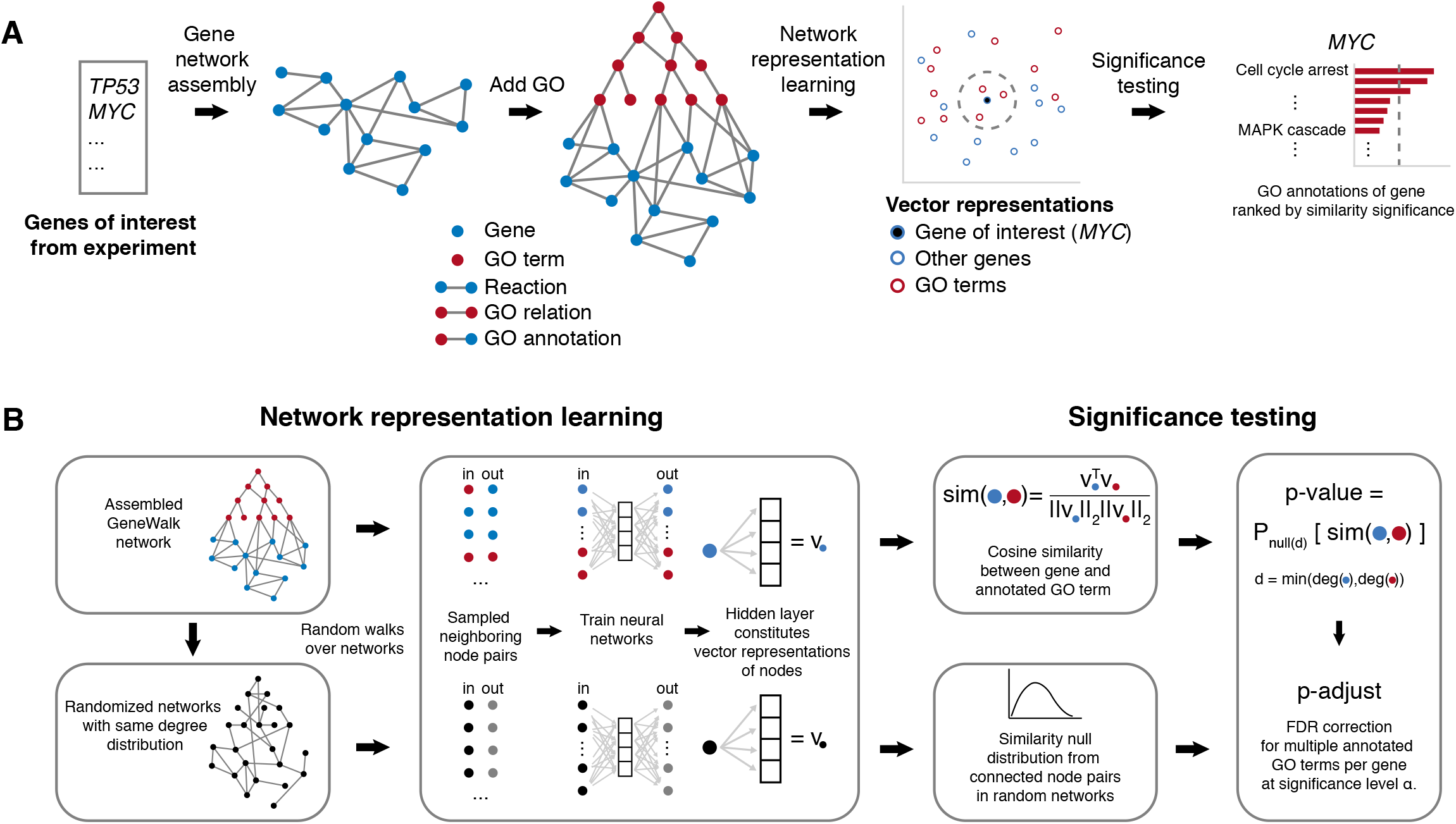
GeneWalk methodology. **A** Schematic introducing the key aspects of the GeneWalk method. The input is a list with genes of interest, e.g. all differentially expressed genes under a certain experimental condition. Using the INDRA^15,16^ or Pathway Commons^14,21^ knowledge base, all molecular reactions in which these genes are involved, are retrieved and assembled in a condition-specific gene regulatory network, to which GO ontology and annotations are then connected. Through network representation learning, each gene and GO term can be represented as vectors, suitable for similarity significance testing. For each gene, GeneWalk gives as output the similarities with all of the connected GO terms, along with their significance, specifying which annotated functions are relevant under the experimental condition. **B** Schematic details of the network representation learning and significance testing parts of GeneWalk. Random walks are generated from the assembled GeneWalk Network (GWN), yielding a large collection of sampled neighboring node pairs, which form the training set of (input, output) of a fully connected shallow neural network (NN) where each node from the GWN is represented as a single feature. The learned hidden layer is the vector representation of a node. And the similarity of a node pair then equals the cosine similarity between the corresponding node vectors. To enable similarity significance testing, we generated randomized networks that were also subjected to DeepWalk and whose resulting cosine similarity values form the null distributions used to calculate a p-value of the experimental similarities between a gene and GO term node. Finally, because multiple GO terms were tested for each gene, we applied an FDR correction (see methods for full methodology).

## Results

### Assembly of a condition-specific GeneWalk Network

The first step in GeneWalk is assembly of a network that describes the relationships between genes and GO terms, starting with a list of relevant genes obtained from a specific experimental assay (Figure 1A). These genes could be differentially expressed (DE) between some condition (such as a genetic mutation or drug treatment) and a control experiment, or the results of a high-throughput genetic screen. A context-specific gene network (Figure 1A) is then assembled using a knowledge base such as INDRA^15,16^. Collections of INDRA statements involving at least two different differentially expressed (DE) genes or a DE gene and GO term are assembled into a gene network such that each gene is represented as a node and each statement as an edge (Figure 1A). This gene network is appended to a GO network^5^ in which edges represent ontological relationships between GO terms as nodes (Figure 1A). To further connect genes to GO terms in the network, we add edges between genes and their annotated GO terms, resulting in a full GeneWalk network (GWN). For comparison, we also generated context-specific gene networks using Pathway Commons^14,21^, which generally had fewer gene-gene and no gene–GO relations.

### Network representation learning with random walks

To determine how genes and GO terms constituting GWN nodes relate to one another, we perform random walks in the network. A network representation learning algorithm (Deep Walk^10^) transforms the random walks into descriptions of how the nodes are embedded in the network, yielding vector representations for each node (Figure 1B). Specifically, short random walks sample the local neighborhood of all nodes, providing a collection of neighboring node pairs, which in turn form a training set of input–output pairs for a neural network (NN) with one hidden layer (Figure 1B). This NN learns which output nodes have been sampled for a given input node from the GWN. After training, the resultant hidden layer weights form the vector representation of any GWN input node (Figure 1B). In this way, groups of interconnected genes and GO terms that are mechanistically or functionally linked to each other occur most frequently as sampled gene–GO term pairs, which can be scored by the cosine similarity between their NN-derived vector representations (Figure 1B).

### Gene-GO term similarity significance testing

Next, GeneWalk calculates whether the similarity values between a gene and GO terms are higher than expected by chance using a significance test (Figure 1B). A null distribution of similarity values between node vectors was generated using representation learning on networks with randomly permuted edges (Supplemental Figure 1A). Comparisons with the null distribution yield p-values for all experimental gene–GO term pairs, which are then corrected for multiple GO annotation testing using the Benjamin-Hochberg false discovery rate (FDR), yielding an adjusted p-value (p-adjust). To decrease variability arising from stochastic walk sampling, network representation learning and significance testing are repeated 10 times to generate the mean and 95% confidence intervals of the p-adjust estimates as the final outputs. The p-adjust values rank the context-specific relevance of all annotated GO terms for a gene of interest. An FDR threshold can then be set to classify annotated GO terms as significantly similar. Gene function significance arises through a high degree of interconnections with other functionally-related genes in the GWN. So genes with many significant functions are likely central to the specific biological context and thus are prime candidates for further investigation.

### Application of GeneWalk to mouse brain RNA-seq data

To test GeneWalk, we applied it to an experimental context in which phenotypes and molecular mechanisms are already well characterized. In the brain (Figure 2A), neurons are myelinated in a *Qki*-dependent manner by oligodendrocytes^22,23^. The *Qki* gene encodes an RNA binding protein involved in alternative splicing^22,23^, and conditional *Qki* deletion in mouse oligodendrocytes (Figure 2A) results in severe hypomyelination and death of the animal^22^. Analysis of RNA-seq comparing animals with *Qki*-deficient and *Qki*-proficient oligodendrocytes^23^ revealed 1899 DE genes (Figure 2B). Several of the strongly downregulated genes had been previously characterized as essential for myelination^24–27^, including *<u>M</u>yelin <u>a</u>nd <u>l</u>ymphocyte protein (Mal), <u>Pl</u>asmo<u>l</u>i<u>p</u>in (Pllp)* and *<u>P</u>roteo<u>l</u>ipid <u>p</u>rotein <u>1</u> (Plp1)* (Figure 2B).

**Figure 2.**
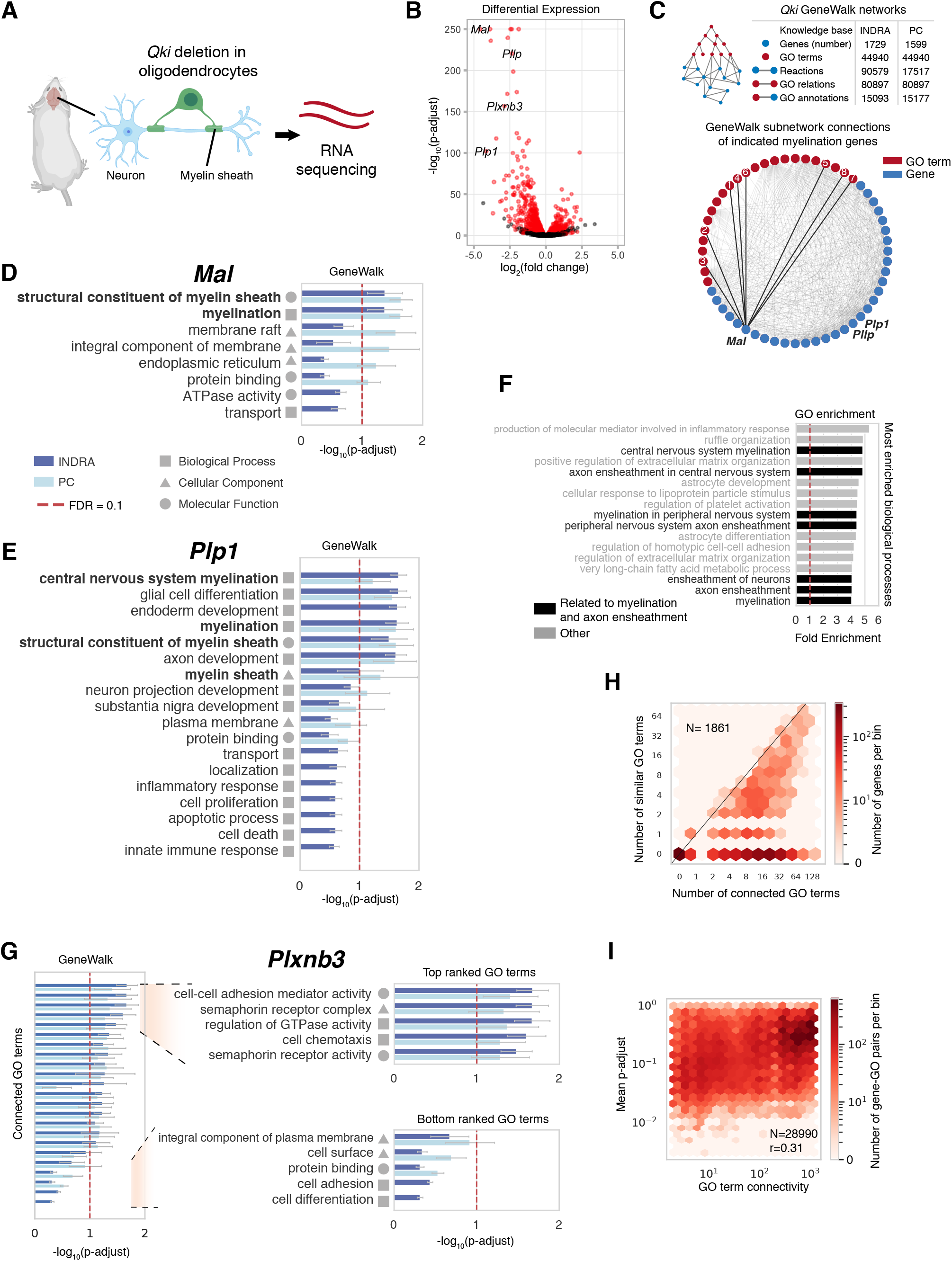
GeneWalk identifies myelination functions from mouse brain RNA-seq. **A** Schematic of the experimental design in Darbelli et al, 2017^23^. Deletion of *Qki*, a gene that encodes RNA-binding proteins, in oligodendrocytes results in hypomyelination in the mouse brain. RNA-seq was performed on Qki’-deficient and control mice (each three biological replicates). **B** Volcano plot showing the results of a differential expression (DE) analysis that compares gene expression profiles in mouse brains with/without *Qki* deletion, re-visualized from^23^. DE genes (N=1899) are indicated in red, which are used as an input to GeneWalk. All other genes are depicted in black. **C** Schematic with statistics of the *Qki*-deficient GeneWalk networks (GWNs) using either INDRA or Pathway Commons (PC) as a knowledge base. Also shown is a visualization of the GWN subnetwork of myelination-related genes *Mal, PllP*, and *Plp1*, all their connected genes and GO terms. Edges (grey) connecting node pairs indicate the presence of INDRA reaction statements or GO annotations between the two respective nodes. Edges between *Mal* and its GO connections (numbered according to rank order in **D**) are highlighted (bold). **D** GeneWalk results for *Mal* in the *Qki*-deficient condition using either INDRA or Pathway Commons (PC) as a knowledge base source to assemble the GeneWalk network. All GO terms connected to *Mal* are rank-ordered by Benjamini-Hochberg FDR-adjusted p-value (p-adjust), indicating their functional relevance to *Mal* in the context of *Qki* deletion in oligodendrocytes. Error bars indicate 95% confidence intervals of mean p-adjust. FDR=0.1 (dashed red line) and domains of GO annotations (square: biological process, triangle: cellular component, and circle: molecular function) are also shown. Supplemental Tables 1 and 2 show full GeneWalk results using the INDRA or PC knowledge base, respectively. **E** As in (**D**) for *Plp1*. **F** Enriched Biological Process GO terms (top 17 shown of 379 enriched terms; Fisher exact test, FDR = 0.01) in *qki*-deficient condition, ranked by fold enrichment, resulting from GO enrichment analysis with PANTHER^1^. Enriched terms related to myelination and axon ensheathment, as reported previously in Darbelli et al, 2017^23^, are highlighted in black. Red line indicates a fold enrichment value of 1, indicating the background. **G** As in (**D**) for *Plxnb3*, the most strongly downregulated DE gene that also has more than half of its connected GO terms classified as significantly similar. The top and bottom ranked GO terms are described in the inset. **H** Hexagon density plot for all genes of interest (N=1861) in terms of number of connected GO terms and number of similar GO terms (at FDR=0.05) resulting from the *Qki*-deficient condition GeneWalk using INDRA as a knowledge base. **I** Hexagon density plot of all tested gene-GO pairs (N= 28990) as a function of GO term connectivity and similarity significance (p-adjust, Pearson correlation r=0.31) for the GWN described in (**H**).

We initiated GeneWalk with 1861 unique Mouse Gene Database (MGD) identifiers^28^ corresponding to the DE gene set (Figure 2B), of which 94% (1750) mapped to different human orthologs using INDRA’s integrated mouse-to-human gene mappings^28,29^. INDRA statements were retrieved for 83% of the genes, of which the vast majority (82% of the initial 1861) had at least one connected GO term (Figure 2C). GeneWalk determined that annotated GO terms related to myelination were most similar to DE genes *Mal, Plp1*, and *Pllp* (Figure 2D-E, S1B), verifying that GeneWalk can identify relevant GO terms for each of these genes. GO enrichment analysis with PANTHER^1^ revealed that myelination-related biological processes were enriched among the DE gene set (Figure 2F, Fisher Exact test, FDR = 0.01)^23^. However, it remains difficult to retrieve gene specific information in this way. Focusing on *Mal* as an example, we find that it is absent from the gene sets corresponding to these most highly enriched GO terms and only first appears as part of the 15^th^ and 17^th^ term “ensheathment of neurons” (108 genes) and “myelination” (106 genes) respectively (Figure 2F). By contrast, GeneWalk associates *Mal* with its direct annotations “myelination” and “structural constituent of myelin sheath” (Figure 2D), a molecular function GO term that was not enriched (p-adjust > 0.05). GeneWalk therefore recovers the known molecular functions of *Mal* in oligodendrocytes whereas enrichment analysis does not.

We assessed the overall performance of GeneWalk: 27% (499) of the DE genes in the GWN had at least one similar GO term (mean p-adjust < 0.05, Supplemental Table 1). The total number of similar GO terms was smaller than expected by chance (KS test, p < 10^−16^; Supplemental Figure 1C), indicating that GeneWalk is selective by focusing on the statistically relevant genes and their functions. Under these conditions, results from GeneWalk were reproducible: no significant differences were observed for the mean p-adjust values when GeneWalk was re-run (two-tailed t-test with FDR correction: all adjusted p-values equal 1.0). For comparison, we also performed a GeneWalk analysis using Pathway Commons, which provided 5-fold fewer reaction statements (Figure 2C) compared to the INDRA knowledge base. INDRA also provides gene–GO term connections obtained from the literature, for example *Plp1* and “inflammatory response” (Figure 2E), whilst GeneWalk with Pathway Commons utilizes GO annotations only. The ordering of GO term significance for these myelination genes was similar regardless of whether Pathway Commons or INDRA was used to generate the GWN (Figure 2D-E,G, Supplemental Figure 1B), demonstrating that GeneWalk is robust to differences in the underlying knowledge base and the amount of available molecular information. These results demonstrate that GeneWalk successfully detects condition-specific gene functions relevant to the biological context of hypomyelination in the mouse brain.

### Generation of gene-specific functions and systematic hypotheses for *Plxnb3* using GeneWalk

We next examined DE genes whose context-specific functions were not clear from previous work. *Plxnb3* was strongly downregulated upon *Qki* deletion (Figure 2B). GeneWalk revealed that more than half of its connected GO terms were significantly similar (mean p-adjust < 0.1), indicating that many of its annotated functions were affected by the *Qki* deletion (Figure 2G). *Plxnb3* is expressed in oligodendrocytes specifically^30^, but it is not annotated to be involved in myelination or related to *Qki* (Figure 2G, Supplemental Table 1). Furthermore, a Pubmed search of *Plxnb3* with the query terms “myelination” or *“Qki”* yielded no results. The most similar functions of *Plxnb3* were “cell–cell adhesion mediator activity”, “semaphorin receptor complex”, “regulation of GTPase activity”, “cell chemotaxis” and “semaphorin receptor activity” (Figure 2G), suggesting that *Plxnb3* could contribute to the myelination process through one of these activities. With GO enrichment analysis, *Plxnb3* was not an element of the gene sets corresponding to the most enriched biological processes (Figure 2G). “Semaphorin–plexin signaling pathway” came 565th when ranked by p-adjust from its Fisher exact test and 73^rd^ by fold enrichment (3.2, p-adjust = 0.03). These results illustrate how GeneWalk determines gene-specific functions and novel hypotheses in a systematic manner and highlights genes that may be overlooked with current methods.

### GeneWalk determines function relevance independent of the degree of annotation

Genes are annotated with different numbers of GO terms. To determine whether GeneWalk is biased with respect to the number of connected GO terms per gene node (the annotation degree), we compared the number of significant GO terms to node degree. The annotation degree is known to introduce a bias into enrichment analyses based on the Fisher Exact test, which overestimates significance for GO terms with large annotated gene sets^8^. We found that with GeneWalk, the distribution of similar GO terms was relatively uniform for all DE genes (Figure 2H, Likelihood Ratio test, χ^2^-test p-value=1), showing that there was no correlation between the numbers of connected and similar GO terms. When we considered only gene-GO term connections originating from INDRA through its automated literature reading functionality, as opposed to GO annotation, we also observed a dispersed distribution (Supplemental Figure 1D), although it was not completely uniform (Likelihood Ratio test, χ^2^-test p-value < 10^−16^). The results show that GeneWalk does not suffer from biases in significance testing towards genes with high or low degrees of annotation.

We also asked whether a GO term with high connectivity is more likely to exhibit strong similarity to a gene simply because it is a highly connected node in the GWN. We found that this was not the case in general (Figure 2I), although there was a weak correlation between the number of connections for a GO term and GeneWalk p-adjust values (Pearson correlation coefficient r=0.31). This effect could mostly be explained by a few highly connected GO terms (Figure S1E), e.g., “cell proliferation” (1152 connections), “apoptotic process” (967 connections), or “localization” (536 connections), for which INDRA detects many genetic associations reported in the literature. However, these GO terms reflect high-level biological concepts that are rarely the specific function of an individual gene. Indeed, in the Pathway Commons-derived GWN, which only contains GO annotations, these GO terms have far fewer connections (42, 33 and 12, respectively), and the correlation between connectivity and similarity significance was lower (r=0.15; Supplemental Figure 1F). Therefore, we conclude that GeneWalk controls for concept generality in GO term relevance ranking and does not suffer from substantial biases related to the degree of GO term connectivity.

### Nascent transcriptome response to bromodomain inhibitor JQ1 using human NET-seq

To test GeneWalk in a different setting, we reanalyzed published NET-seq data^31^ describing the response of a human T-cell acute lymphoblastic leukemia cell line (MOLT4) to treatment with JQ1 (Figure 3A), a small molecule that targets the BET bromodomain in BRD4 and other BET family members^32^. NET-seq measures RNA polymerase position genome-wide at single-nucleotide resolution^31,33^, yielding a quantitative description of the nascent transcriptome. We calculated Pol II coverage per gene and identified differentially transcribed protein-coding genes using DEseq2^2^ (Figure 3B). INDRA statements were retrieved for 82% of DE genes (N=2670), 79% of which had connected GO terms. GeneWalk identified similar GO terms for 28% of DE genes (mean p-adjust < 0. 05, Supplemental Table 3), similar to the statistics for the mouse brain RNA-seq data. Conventional GO enrichment analysis of the JQ1 DE gene set only yielded five high-level (generic) functions such as “ncRNA metabolic process” and “chromatin organization” with low fold enrichment (range, 1. 2-1.7; Figure 3C, Fisher exact test, FDR = 0.05). These results reveal the limitations of GO enrichment analysis when many functionally unrelated genes are misregulated. GeneWalk does not suffer from this limitation, because it is based on the local regulatory network connectivity with other treatment-affected genes.

**Figure 3.**
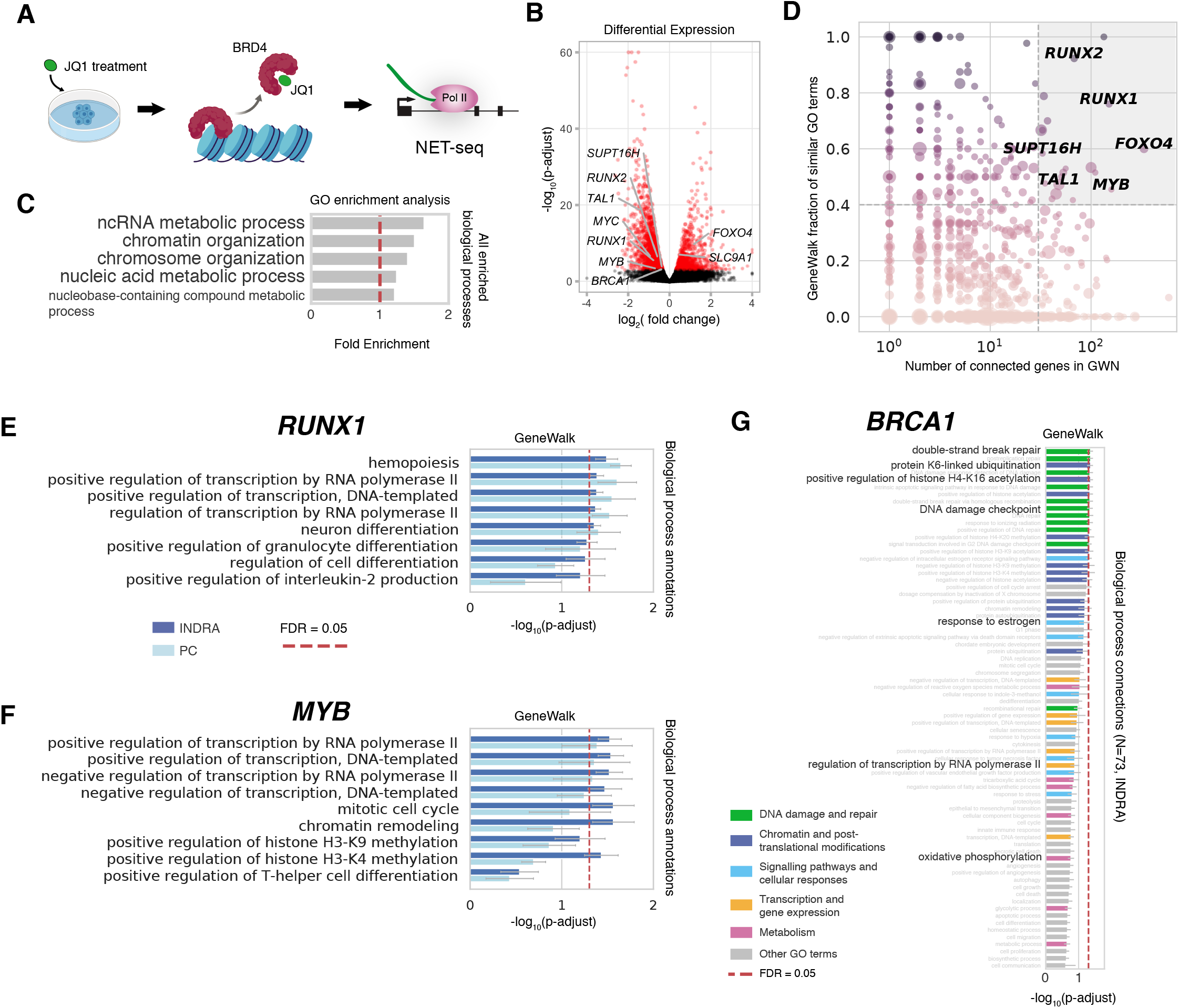
GeneWalk analysis of nascent transcriptome response to BRD4 inhibition in T-ALL cells. **A** Schematic of the experimental design in Winter et al, 2017^31^. NET-seq was performed on JQ1-treated MOLT4 cells (1 μM for 2 hours, alongside DMSO controls, two biological replicates each). JQ1 targets BRD4 and other BET bromodomain family members, causing BRD4 to dissociate from chromatin^31^. **B** Volcano plot showing the results of a differential expression (DE) analysis comparing RNA Polymerase II gene coverage between JQ1 and DMSO control samples. DE genes (N=2692), indicated in red, were used as an input to GeneWalk. All other genes are depicted in black. **C** All enriched Biological Process GO terms (five enriched terms, Fisher exact test, FDR = 0.05) in JQ1 condition, ranked by fold enrichment, obtained by GO enrichment analysis using PANTHER^1^. Red line indicates a fold enrichment value of 1, indicating the background. **D** Scatter plot with DE genes as data points showing the GeneWalk fraction of similar GO terms over total number of connected GO terms (min_f, minimum value between INDRA and PC GWNs) as a function of the number of gene connections in the GWN (N^gene^, again minimal value between INDRA and PC). The circle size scales with the differential expression significance strength (-log_10_(p-adjust)) and the color hue with min_f. Twenty genes were identified with min_f > 0.4 and N^gene^ > 30 (grey-shaded area, see Table 1 for complete list). **E** GeneWalk results for the transcriptional regulator *RUNX1* under JQ1 treatment. Annotated biological process terms are rank-ordered by mean FDR adjusted p-value. Error bars indicate 95 % confidence intervals of mean p-adjust. FDR=0.05 (dashed red line) is also shown. See Supplemental Table 3 (INDRA) and 4 (PC) for full details. **F-G** As in (**D**) for transcriptional regulator *MYB* (**F**) and *BRCA1* (**G**). For *BRCA1*, INDRA annotations are indicated by class: DNA damage and repair (green), Chromatin and post-translational modifications (dark blue), signalling pathways and cellular responses (light blue), transcription and gene expression (yellow), metabolism (purple) an other GO terms (grey).

### GeneWalk identifies transcriptional regulators responding to JQ1 treatment

To identify genes that were highly affected by JQ1 treatment we focused on those with a high fraction of similar GO terms over all connected terms according to GeneWalk with both INDRA and Pathway Commons knowledge bases (Figure 3D, fraction > 0.4 and DE p-adjust < 0.01). We reasoned that by further selecting for genes with a large connectivity with other DE genes (Figure 3D, gene connectivity > 30), we might identify candidate genes that mediate the observed transcriptional changes. With this procedure, we identified 20 genes (Figure 3D, Table 1), of which 13 (Fisher Exact test, odds ratio=12, p=2×10^−7^) had significantly similar transcription-related annotations (Table 1). When also including gene - GO term relations obtained through the literature with INDRA, this number rose to 16 (Fisher Exact test, odds ratio=10 p=2×10^−6^, Table 1). Among these were *RUNX1* (Figure 3E), *MYB* (Figure 3F), *RUNX2*, and *TAL1* all genes that have previously been identified as part of a core transcriptional circuitry important to our leukemia model system^31^. We also found newly implicated genes such as *SUPT16H* (Figure S2A), with its most similar cellular component term being “FACT complex” (mean p-adjust = 0.01, Supplemental Table 4), as expected, and *FOXO4* (Figure S2B) with relevant molecular functions such as “RNA polymerase II transcription factor activity, sequence-specific DNA binding” (mean p-adjust=0.03, Supplemental Table 4). These results demonstrate the capability of GeneWalk to systematically identify genes with relevant context-specific functions.

**Table 1.** GeneWalk identifies transcriptional regulators among highly connected genes with many significant functions in the JQ1 condition.

| Gene | Most similar biological process annotation<br>(GeneWalk with Pathway Commons knowledge base) | Mean<br>p-adjust | Significant<br>transcription<br>annotation<br>(FDR=0.05)? |
| --- | --- | --- | --- |
| *RUNX1* | Hemopoiesis | 0.023 | Yes |
| *RUNX2* | Hemopoiesis | 0.019 | Yes |
| *MYB* | Positive regulation of transcription by RNA polymerase II | 0.041 | Yes |
| *TAL1* | Positive regulation of transcription by RNA polymerase II | 0.032 | Yes |
| *SUPT16H* | DNA replication-independent nucleosome organization | 0.014 | Yes |
| *FOXO4* | Positive regulation of transcription by RNA polymerase II | 0.033 | Yes |
| *RBBP8* | DNA double-strand break processing involved in repair via single-strand annealing | 0.013 | Yes |
| *RREB1* | Positive regulation of transcription by RNA polymerase II | 0.013 | Yes |
| *TFAP4* | Positive regulation of transcription | 0.027 | Yes |
| *HIST2H2AC* | Chromatin organization | 0.010 | No |
| *GABPB2* | Positive regulation of transcription by RNA polymerase II | 0.032 | Yes |
| *TP53BP1* | Double-strand break repair via nonhomologous end joining | 0.018 | Yes |
| *ELOVL6* | Fatty acid elongation, saturated fatty acid | 0.024 | INDRA only |
| *DICER1* | Conversion of ds siRNA to ss siRNA involved in RNA interference | 0.024 | No |
| *CDC5L* | Positive regulation of transcription by RNA polymerase II | 0.058 | Yes |
| *CDKN1A* | DNA damage response, signal transduction by p53 class mediator resulting in cell cycle arrest | 0.034 | INDRA only |
| *CUL1* | SCF-dependent proteasomal ubiquitin-dependent protein catabolic process | 0.010 | No |
| *EPAS1* | Cellular response to hypoxia | 0.021 | Yes |
| *EDNRA* | Artery smooth muscle contraction | 0.039 | INDRA only |
| *SLC9A3* | Sodium ion import across plasma membrane | 0.016 | No |

### GeneWalk quantitatively ranks GO annotation relevance for genes encoding moonlighting proteins

Genes with a large number of GO annotations encode proteins that are involved in multiple molecular functions or biological processes and thus could be moonlighting proteins^34^. As these functions might not all be relevant in a particular context, GeneWalk is well suited to identify the relevant functions for genes encoding moonlighting proteins. We identified 9 DE genes with at least 30 connected GO terms, of which no more than 40% were significantly similar (Figure S2C). Among them was *MYC*, a widely studied proto-oncogene and member of the reported core transcriptional circuitry^31^. But *MYC* was not identified with our transcriptional regulator analysis (Figure 3C), because GeneWalk determined that its majority of annotations, most of them unrelated to transcription, were insignificant in the JQ1 condition (Supplemental Table 4). *BRCA1* was another downregulated gene (Figure 3B, S2C) with 23% (17) of its 73 connected biological processes being significantly similar (Figure 3G, FDR=0.05, Supplemental Table 3). GeneWalk ranked DNA damage and repair-related processes as most relevant (Figure 3G, mean p-adjust < 0.05), followed by histone and other post-translational modification-related terms (mean p-adjust = 0.05-0.07). Transcription, metabolism and other GO terms were the least relevant (mean p-adjust > 0.09). These results demonstrate the capability of GeneWalk to systematically prioritize context-specific functions over less plausible alternatives, which is especially useful when considering genes encoding moonlighting proteins.

### GeneWalk facilitates functional characterization of cellular response to isoginkgetin

As a final application of GeneWalk, we studied the transcriptional response to the biflavonoid isoginkgetin (IsoG), a plant natural product and putative anti-tumor compound whose mechanism of action remains unknown. IsoG inhibits pre-mRNA splicing *in vitro* and *in vivo*^35^ and also causes widespread accumulation of PoI II at the 5’ ends of genes, indicating an additional role as a Pol II elongation inhibitor^36^. Through DE analysis of NET-seq data obtained from HeLa S3 cells treated with IsoG (Figure 4A), we identified a total of 2940 genes as differentially transcribed, most of which exhibited upregulated Pol II occupancy (Figure 4B, FDR=0.001). Using INDRA and Pathway Commons as the knowledge bases, we applied GeneWalk to these DE genes and found that 24% had at least one similar GO term (FDR = 0.05, Supplemental Table 5).

**Figure 4.**
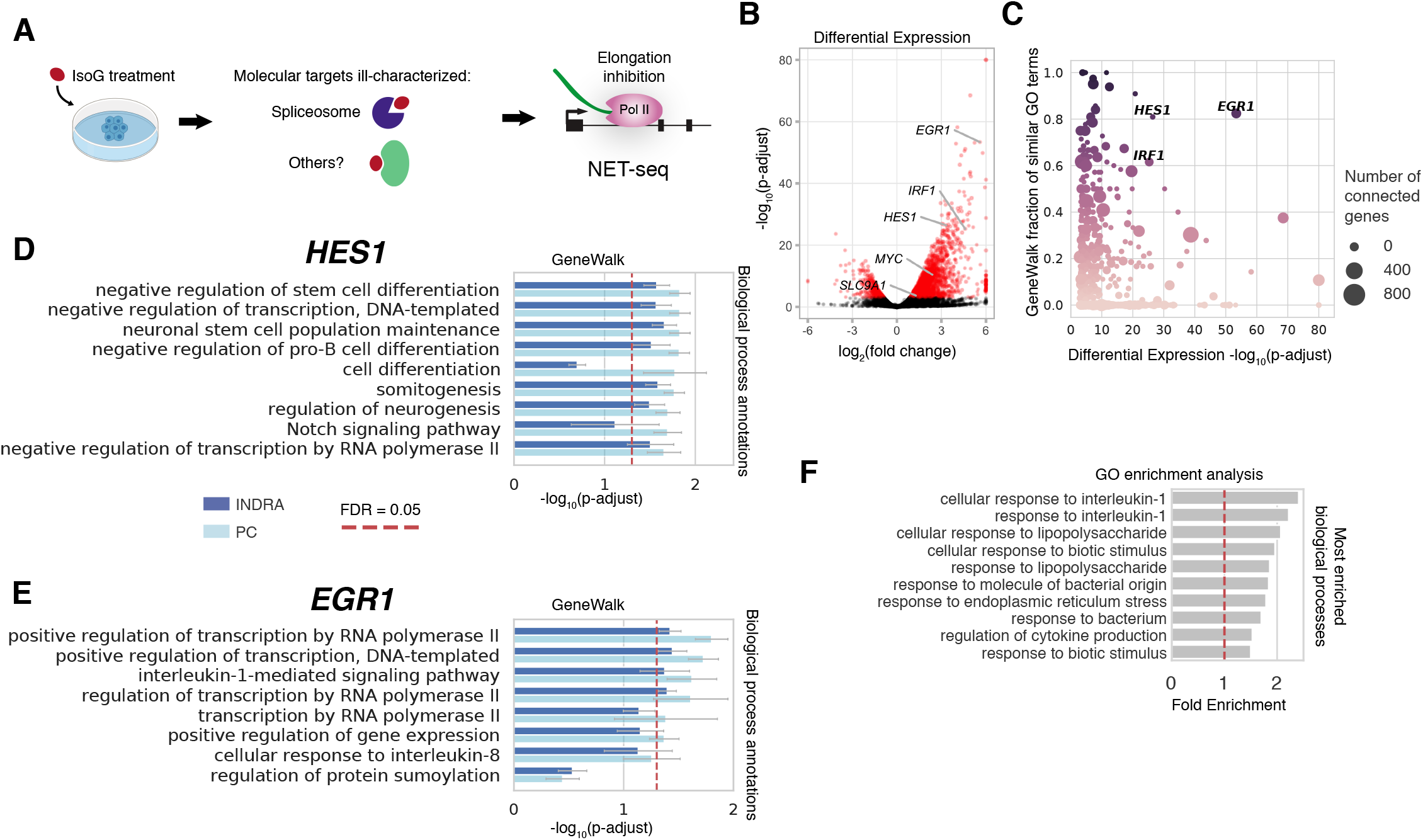
GeneWalk determines condition-specific functions of regulators through comparison of nascent transcriptome response to IsoG and JQ1 treatment. **A** Schematic of the experimental design in Boswell et al, 2017^36^. NET-seq was performed on isoginkgetin (IsoG)-treated HeLa S3 cells (30 μM IsoG for 6 hours, alongside DMSO controls, two biological replicates each). The *in vivo* molecular targets remain incompletely characterized, as IsoG treatment causes widespread Pol II elongation inhibition. **B** Volcano plot showing the results of a differential expression (DE) analysis that compares RNA Polymerase II gene coverage between IsoG and DMSO control samples. DE genes (N=2980), indicated in red, were used as an input to GeneWalk. All other genes are depicted in black. **C** Scatter plot with DE genes as data points showing the fraction of similar GO terms over total number of connected GO terms (min_f, minimum value between INDRA and PC GWNs) as a function of the DE significance strength (-log_10_(p-adjust)). The circle size scales with number of connected genes in GWN (INDRA) and the color hue with min_f. **D** GeneWalk results for *HES1* under IsoG treatment with INDRA or PC knowledge bases. Annotated biological process terms are rank-ordered by mean FDR adjusted p-value (p-adjust). Error bars indicate 95% confidence intervals of the p-adjust estimate. FDR=0.05 (dashed red line) is also shown. See Supplemental Table 5 (INDRA) and 6 (PC) for full details. **E** As in (**D**) for *EGR1*. **F** Enriched Biological Process GO terms (top 10 shown of 41 enriched terms; Fisher exact test, FDR = 0.01) in the IsoG-treated condition, ranked by fold enrichment, obtained by GO enrichment analysis using PANTHER^1^. Red line indicates a fold enrichment value of 1, indicating the background.

To gain insight into the molecular mechanisms of IsoG-induced transcriptional changes, we searched for genes that were both strongly differentially expressed (p-adjust < 10^−25^) and had a large fraction of functions significantly affected according to the GeneWalk analyses with both INDRA and Pathway Commons (Figure 4C, fraction > 0.5). In this manner, we identified three genes: *HES1, EGR1* and *IRF1* (Figure 4C). *HES1* had “negative regulation of transcription, DNA templated” as one of the most similar biological processes (Figure 4D, S3A) and has been reported to inhibit transcription elongation^37^. *EGR1* and *IRF1* both had as most similar term “positive regulation of transcription by RNA polymerase II” (Figure 4E, S3B). Based on our GeneWalk analysis, we propose the hypothesis that *EGR1* and *IRF1* could contribute to transcriptional initiation upregulation whereas *HES1* would be the lead candidate to cause Pol II pausing during transcription elongation.

By contrast, GO enrichment analysis of IsoG treatment revealed 41 enriched terms (FDR = 0.01), of which 10 had a fold enrichment larger than 1.5 (Figure 4F), including “cellular response to interleukin-1” and “cellular response to biotic stimulus”. Also, “inflammatory response” was weakly enriched (fold enrichment = 1.6, p-adjust = 0.05). Indeed, interleukins such as *IL1A, CXCL2, CXCL3* and *CXCL8* were upregulated and had similar GO annotations related to inflammatory response or innate immune response according to GeneWalk (Supplemental Table 5-6, mean p-adjust < 0.05). However, no terms related to transcription were enriched other than the generic term “positive regulation of gene expression” (fold enrichment = 1.2, p-adjust = 0.05) with a corresponding gene set that includes 379 DE genes. According to GeneWalk, *EGR1* also had “interleukin-1-mediated signaling pathway” as a similar function (Figure 4E), providing a possible link between the interleukins and observed transcriptional changes. We conclude that GO enrichment analysis results were consistent with those from GeneWalk. However, GO enrichment failed to provide insight for the molecular mechanism of IsoG-induced transcriptional changes, whereas GeneWalk was able to provide *HES1* and *EGR1* as plausible candidate genes given the available body of knowledge.

### Comparison between JQ1 and IsoG analyses indicates that GeneWalk yields condition-specific gene functions

To confirm that GeneWalk’s function assignments are not constant and depend on the experimental condition, we compared GeneWalk analyses of JQ1 and IsoG treatments. Between the JQ1 and IsoG condition, 538 DE genes were shared (Figure 5A), marginally more than expected by chance (Fisher Exact test: p=0.02, odds ratio = 1.1, 95% confidence interval: [1.0, 1.3]). As examples, we compared the overlap of similar GO terms of *MYC* and *SLC9A1*, which are common DE genes between JQ1 (Figure 3B) and IsoG treatment (Figure 4B). *MYC* is annotated to be involved in 29 biological processes (Figure 5B). Between the two GeneWalk analyses, *MYC* showed 5 significant biological processes and 9 molecular functions for IsoG and 0 and 1 respectively for JQ1 (Figure 5B,S3C, FDR = 0.05). “Nucleus” and “nucleoplasm” were significant cellular components in both conditions (Figure S3D). For *SLC9A1*, different biological processes were significant for each condition. For example, *SLC9A1* had “response to acidic pH” as relevant only for the JQ1 context and “cellular sodium ion homeostasis” specific to IsoG treatment (Figure 5C, FDR = 0.05).

**Figure 5.**
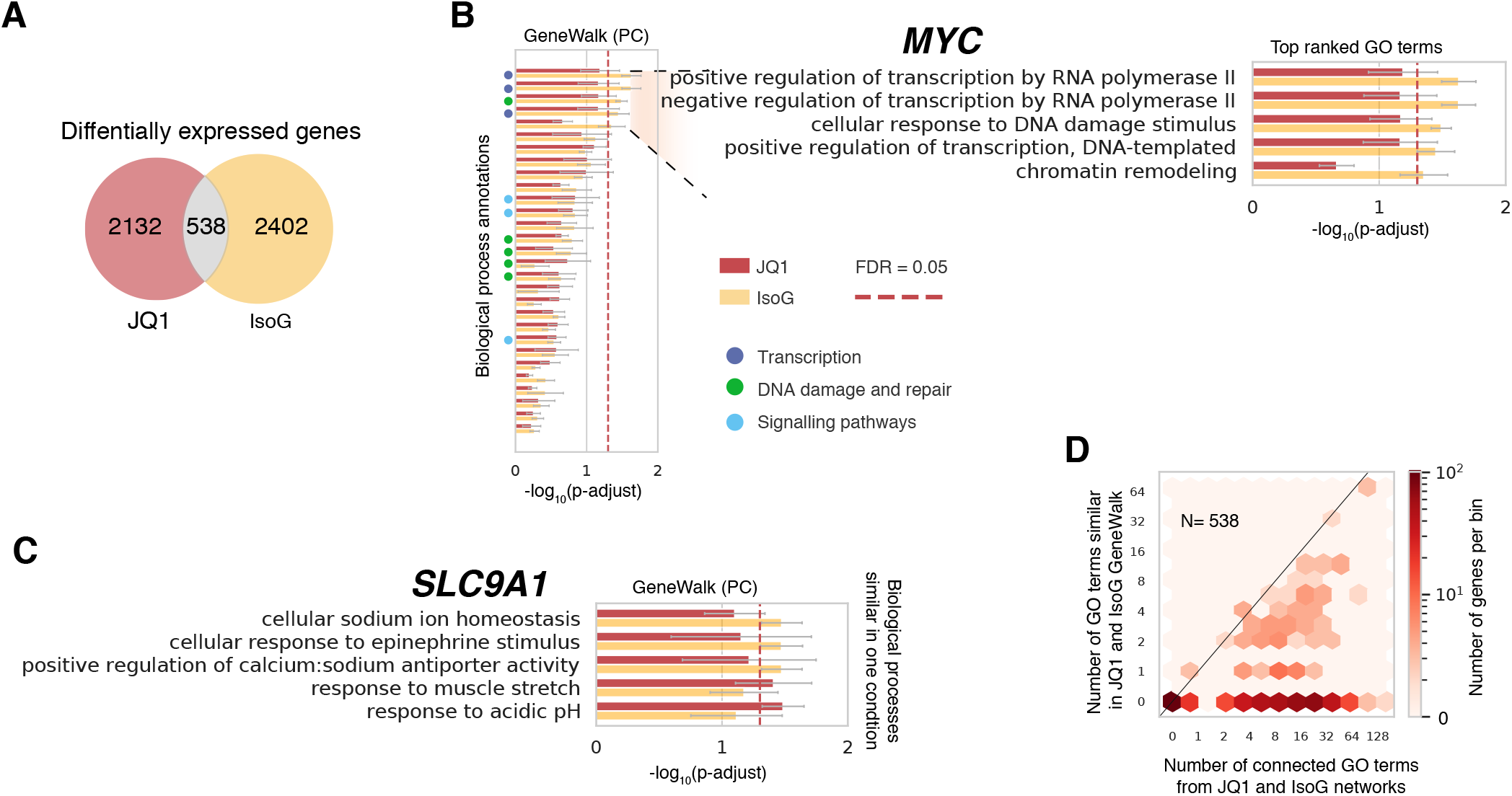
JQ1 and IsoG functional overlap analysis indicates GeneWalk determines context-specific functions. **A** Venn diagram detailing the overlap (Fisher exact test: p=0.02, odds ratio = 1.1, 95% Confidence Interval: [1.0, 1.3]) of DE genes between JQ1 and IsoG treatments as described in Figure 3B and 4B. **B** GeneWalk results (with PC as data source) for *MYC* in the JQ1 (red) and IsoG (yellow) condition. Annotated biological processes are rank-ordered by FDR-adjusted p-value, indicating the relative functional importance of transcription (dark blue), DNA damage and repair (green), and signalling pathways (light blue) to *MYC* under the IsoG condition. The top five most similar GO terms are described in the insets. See Supplemental Table 4 (JQ1) and 6 (IsoG) for full details. Red dashed line indicates FDR=0.05. **C** As in (**B**) for *SLC9A1*, showing the biological process terms that are significantly similar in either JQ1 or IsoG condition. **D** Hexagon density plot for overlapping DE genes (N=538) in terms of number of overlapping similar GO terms and number of possible shared connected GO terms for the GeneWalk network using INDRA as a knowledge base.

Overall, the numbers of shared similar GO terms determined by GeneWalk were relatively uniformly distributed (Figure 5D, Likelihood Ratio test, χ^2^-test p-value=1), indicating a lack of systematic bias in function assignment. Many genes had no shared terms between the two drug treatments (Figure 5D), suggesting that those DE genes have different roles in each condition. We found similar results for GO terms originating from INDRA (Supplemental Figure S3E, Likelihood Ratio test, χ^2^-test p-value=1). We conclude that GeneWalk is able to determine context-specific functions as a consequence of differences in the context-specific gene-gene interactions part of the GeneWalk network.

## Discussion

Here we have described GeneWalk, a machine learning and statistical modelling method that identifies condition-specific functions of individual genes. Although we demonstrate its capabilities with differentially expressed genes obtained by two experimental approaches, RNA-seq and NET-seq, we emphasize that GeneWalk is capable of analyzing gene hit lists arising from many other types of experimental assays, such as CRISPR screens or mass spectrometry. In principle, for any gene of interest it is possible to recover relevant information by manual searches of the scientific literature. However, manual searching is time consuming when dozens or more genes are involved and potentially biased because manual searches are typically incomplete. In contrast, GeneWalk provides a principled way to score gene-GO annotation associations based on systematic assembly of prior knowledge curated from the scientific literature. Information about context-specific gene functions can lead to hypotheses about gene regulation even when transcriptome-wide enrichment methods fail to yield significant results. If no molecular information on a gene has been reported, GeneWalk does not make any functional relevance prediction. However, this bias towards studied genes is clearly also present for manual searches or enrichment analyses. Currently, only connected GO terms are considered for identification of function relevance, but we imagine that GeneWalk could be extended to predict novel gene functions because of high similarity scores between a gene and unconnected GO terms.

The GeneWalk applications in this study used the INDRA^15,16^ and Pathway Commons^14^ knowledge bases which enable automated assembly of a GeneWalk network. Although these databases are optimized for human genes, we show that when mouse genes can be mapped unambiguously to their human orthologues, a network can still be assembled. For more distant species, this approach is likely to be insufficient. Nevertheless, GeneWalk should be readily applicable in other model organisms, such as yeast, given the availability of annotated gene regulatory networks and GeneWalk’s option to analyze user-provided pre-assembled networks. In summary, we provide GeneWalk as a general open source tool (github.com/churchmanlab/genewalk) for the scientific community to facilitate experiment interpretation and hypothesis generation through systematic and unbiased prioritization of functional genomics experiments conditioned on the largest possible assembly of prior knowledge.

## Methods

### Assembly of mechanistic networks using INDRA

We used the Integrated Network and Dynamical Reasoning Assembler (INDRA) system^15^ to collect and assemble a set of statements from the scientific literature and pathway databases. INDRA integrates content from i) multiple natural language processing systems (REACH^38^ and Sparser^39^) of primary literature in the minable NCBI corpus and ii) queries on pathway databases (Pathway Commons^14,21^, BEL Large Corpus^40^, SIGNOR^41^). INDRA extracts information about molecular mechanisms from these sources in a common statement representation, which has a rich functional semantic with respect to reactant and reaction types. Each statement represents a mechanistic relationship (e.g., activation/inhibition, regulation of amount, or post-translational modification) between two entities or between an entity and a biological process. For each data set described in this study, we queried the pathway databases and machine reading results from REACH and Sparser (run on all Medline abstracts and PubMedCentral manuscripts) for interactions involving the DE genes in the dataset. The resultant set of statements consisted only of relationships among DE genes, GO terms, and protein families and complexes containing the DE genes, obtained from the FamPlex ontology^16^. The final set of statements was then used as an input to the core GeneWalk algorithm as described below.

### Assembly of GeneWalk Network with gene regulation, GO ontology and annotation

To generate the gene network from each context-specific set of INDRA statements, we initialized a networkx (v2.2) multigraph in Python (v3.6) and defined from all statements with at least two different agents (human DE genes, their gene family names and/or GO identifiers), nodes for each agent and edge for the reaction itself (with edge label the reaction type). We added edges (label: ‘FPLX:is_a’) between genes and (if already present in the network) any corresponding gene family names according to relations defined with FamPlex^16^.

When using Pathway Commons (PC) as a source for the gene reactions, we downloaded a simple interaction format (NodeA <relation_type> NodeB) PC network (PathwayCommons11.All.hgnc.sif.gz) from pathwaycommons.org. loaded the PC network as a networkx multigraph (with edge label the relation type), and maintained only the subnetwork of nodes (and existing edges between them) that corresponded to human DE gene symbols. When using a mouse DE gene list as an input, the MGD identifiers are first mapped to their human ortholog HGNC identifiers and gene symbols with INDRA’s integrated HGNC and MGD mappings^28,29^ before proceeding with the network assembly steps described above.

Next, for each gene in the network (originating from either INDRA or PC), we added GO nodes and edges (label: ‘GO:annotation’) for each GO annotation as imported with GOAtools^5^ (v0.8.12) by matching the gene’s UniProt identifier, an attribute provided by INDRA. We only included annotations without a ‘NOT’ qualifier and based on manually reviewed, possibly phylogenetically inferred experimental evidence, i.e. those with the following GO evidence codes^7^: EXP, IDA, IPI, IMP, IGI, IEP, HTP, HDA, HMP, HGI, HEP, IBA, and IBD. Finally, we imported the GO ontologies (Biological Processes, Molecular Function, and Cellular Component, release 2018-06-20), again using GOAtools, and added to the network the remainder of GO term identifiers as nodes and parent relations from the ontology as edges (label: “GO:is_a’). For generality, we also provide a network assembly option: “edge_list,” which allows the user to provide a predefined GWN in an edge list format (text file in which each line indicates an edge connecting respective node pairs: NodeA NodeB), or “sif” (simple interaction format, as mentioned above). It is assumed that the nodes are either gene symbols or GO identifiers.

### Network representation learning using random walks

To learn the vector representations of all nodes in the GWN, we implemented a version of the unsupervised machine learning algorithm DeepWalk^10^ in Python (v3.6) with all hyperparameters optimized to ensure the functionality and reproducibility of GeneWalk. The existence of difference types of evidence can generate multiple edges between a node pair. In order to generate a network that reflects the unique nature of molecular interactions, we collapse such multiple edges, thereby reducing the network from a multigraph to a graph. Thus, for our purposes, the degree d(n) of a node n represents the number of nodes connected by at least one edge in the multigraph. We then sample random walks over the network. A random walk over a network represents a random sequence of nodes that are each directly connected by an edge. The probability p to jump from node n to any connected node equals p=1/d(n). To sample the local neighborhood of a node n_1_, we start in n_1_ and sample a short random walk of a total of L=10 nodes for d(n_1_) times and perform this procedure for each node in the network. To ensure the reproducibility of the resultant vector representations by having sufficient amounts of sampled walks, we repeat the above procedure N_iteration_ = 100 times. Longer walk lengths were tested (L=100, 200, 400, 800, 1600, 4800) but were found to be unsuitable for querying the local neighborhood of each node due to the high network connectivity of the GWN (Supplemental Fig. 1E). Because the effective node distance traveled for a random walk scales with L^12^, shorter walk lengths would not sufficiently sample the local node neighborhood, and were therefore not considered. Lower numbers of iterations (N_iteration_ =1,2,4 or 8) resulted in irreproducible similarity values due to stochastic sampling variation, whereas greater numbers of iterations (N_iteration_=200) did not alter our results relative to N_iteration_=100.

As described in the main text (Figure 1B), the sampled random walks provide a collection of neighboring node pairs, which in turn form a training set of input–output pairs for a fully connected neural network (NN) with one hidden layer of dimensionality d. The NN input and output layers are one hot encodings of all nodes from the GWN. In practice, and as previously described for DeepWalk^10^, this NN is trained through implementation of the word2vec algorithm^42^ (in our case, with gensim package v3.7.1 with the following options: skip-gram model with k=5 negative sampling words and without downsampling: sample=0 and min_count=0 and window/context size=1, dimension d=8; for further documentation see https://radimrehurek.com/gensim/models/word2vec.html). Intuitively, our sampled random sequences of nodes are analogous to sentences, which are then used for training to convert words (nodes) into vector representations. When the window size in word2vec is set to 1, it only considers directly connected node pairs from random sequences. Formally, the loss objective of the word2vec NN with input word *W_I_* and output word *w_o_* is^42^: 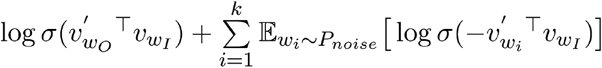, with *P_noise_*(*w*) ∝ *U*{*w*)^3/4^ and *U*(*w*) the unigram distribution. Here, *v^w_I_^* represent the input weights of *w_I_*, which constitute the vector representations used for our GeneWalk analysis, and 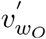 the output weights for *w_O_*. For the vector dimensionality d, we tested different values (2, 3, 4, 6, 8, 12, 16, 32, 50, 500), and found that d=8 was optimal because the variance of the resulting cosine similarity distributions was largest, indicating the highest sensitivity of detection of similarity between node pairs. Lower dimensionality generally resulted in high similarity between all nodes, whereas higher dimensionality lowered all similarity values; both cases resulted in a reduced variability. After training, for any input node from the GWN, the resultant hidden layer weights form the vector representation of that node (Figure 1B). In practice, the gensim package provides a dictionary (gensim.models.word2vec.wv) with all the resultant node vectors, which can then be used for significance testing as described below.

### Determining statistical significance of GeneWalk similarity values

For similarity significance testing, we first generated a randomized network from the GWN, i.e., a network with the same number of nodes as in the GWN but with edges randomly permuted such that the GWN degree distribution is retained (networkx v2.2 configuration_model function). With this random network, we proceed with network representation learning as described above for the GWN to generate random node vectors, which are then used to form null distributions of cosine similarity values (gensim wv.similarity function). In the random networks, node pairs with a low degree of connectivity tend to have lower similarity values than highly connected node pairs (Supplemental Figure 1A). We generated degree-dependent null distributions as follows. For each node n in the random network, we calculate the cosine similarities with all it neighbors to form a null distribution. We repeat this for 10 independently randomized networks and collate the similarity values from all replicates to assemble a sufficiently large degree-dependent distribution. Next, we proceed with significance testing for each connected gene-GO term pair present in the GWN. The p-value for such a pair then equals the normalized rank of the cosine similarity in the null distribution. To correct for multiple GO annotation testing for a given gene, we utilized the Benjamini–Hochberg FDR adjusted p-value (Python package: statsmodels, function: stats.multitest.fdrcorrection). Finally, we repeat the above described network representation learning and significance testing procedures of the GWN 10 times and provide the mean and 95% confidence intervals of p-adjust as our final outputs alongside the mean and standard errors (s.e.m.) of the generated gene-GO pair similarity values.

### Differential expression analysis of mouse RNA-seq

Mouse *Qki* deletion RNA-seq experiments and DE analysis were described previously^23^. The DE results are re-visualized in Figure 2B for completeness.

### Differential expression analysis of NET-seq

JQ1 and IsoG NET-seq experiments were previously described in^31,36^, respectively, and the data are available in GEO accession number GSE79290 and GSE86857. In brief, MOLT4 cells (two biological replicates per condition) were treated either with JQ1 (1 μM, 2h treatment) or DMSO (negative control). For the IsoG study, HeLa S3 cells (two biological replicates per condition) were treated with IsoG (30 μM for 6h) or DMSO control. NET-seq, 2 replicates, DMSO control.

We generated NET-seq coverage files^33^ with modifications described below. Here, we used NET-seq alignment scripts are available at www.github.com/churchmanlab/MiMB2019NETseq. Briefly, we utilized 5’ random hexamers as UMIs by displacing them from the.fastq read sequence and aligned the resultant reads with STAR (v2.5.1a) with genome assembly GRCh38 and annotation from GRCh38v86. We filtered out multimapping alignments, PCR duplicates, and RT mispriming reads (i.e. cases in which RT priming occurred at a position within the RNA fragment instead of the 3’ end), and splicing intermediates. Finally, we generated NET-seq coverage files (.bedgraph format) at nucleotide resolution with HTseq (v0.9.1) using the whole read length.

The coverage files were imported into R (v3.5.0, packages: GenomicRanges v1.32.3, rtracklayer v1.40.3) to determine gene coverage, i.e., the sum over base-pair counts, using Ensembl gene_ids. We filtered for protein-coding genes (annotation acquired with package biomaRt v2.36.1) with positive coverage, i.e., counts per gene averaged over all conditions > 20. The resultant genes and their counts were then utilized to determine differentially expressed genes with DEseq2^2^ (v1.20, default parameters except as follows: FDR=0.001, betaPrior=false and poscount size factor estimation (JQ1) or total read count as size factor for IsoG). After differential expression, we filtered for genes with an HGNC identifier and gene symbol to ensure INDRA could accept them as an input.

### GO enrichment analysis

To perform GO enrichment analysis, we utilized PANTHER^1^ (www.pantherdb.org with options: Overrepresentation Test Released 20181113, GO biological process complete, GO molecular function complete or GO cellular component complete - release 2018-12-01, Fisher Exact test, FDR correction). For JQ1 and IsoG, the analyzed lists consisted of Ensembl identifiers corresponding to the DE genes that were used as an input to INDRA. The reference list contained Ensembl identifiers of all genes tested in the DEseq2 analyses. For the QKI study, GO enrichment analysis was described previously^23^ and reproduced here as described for JQ1 and IsoG, with the exceptions that MGD identifiers were used for all the DE genes and reference list, and GO ontology release 2019-01-01 was used.

### Likelihood ratio test for uniform distribution of similar GO terms

To assess how the number of similar GO terms relates to the number of connected GO terms (Figure 2H), we developed a likelihood ratio test. Without loss of generality, this test is also applicable to other described cases (Figure 4B, S1D and S3C) where the random variable *Y* described on the y axis (number of similar GO terms in Figure 3F) has the intrinsic dependency *Y* ≤ *X* on a random variable described on the x axis *X* (number of connected GO terms in Figure 2H). First, note that for any discrete joint probability distribution 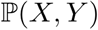, we have a conditional probability relation: 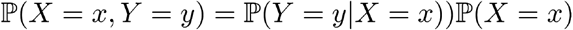. The null hypothesis *H*_0_ for our likelihood ratio test is that *Y|X* is uniformly distributed between 0 and 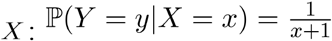. The alternative hypothesis *H*_1_ is that the conditional probability is not uniform, but instead determined by *a priori* unknown probabilities: 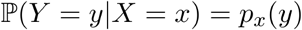.

For any given *x*, if we repeatedly observe *Y|X* — *x* for a total of *N_x_* multiple independent times, the joint frequency function, i.e., the collection of numbers of times {*n_y,x_*} each *V* ∈ {0,1, …, *x*} value is observed, follows a multinomial distribution^43^: 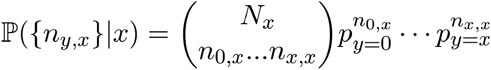, with *P_y_* equal to the above described condition probabilities specific for each hypothesis. The likelihood ratio Λ^43^, is by definition the ratio of joint probabilities functions under each hypothesis with maximum likelihood estimated (MLE) parameter values given our observed data 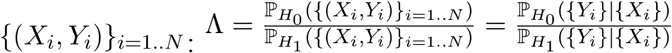. In our case, this is the ratio of multinomial distributions with probabilities defined by each hypothesis. Under *H*_1_, the MLEs equal^43^. 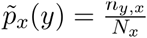 with 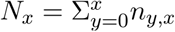 the total number of y observations for a given *x*. On the other hand under *H*_0_, the uniform distribution fully determines the probabilities and are thus independent of our observations: 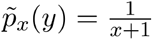. Now let 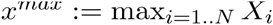 be the maximum observed X value. Thus, our likelihood ratio reduces to: 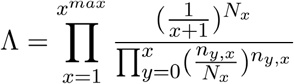.

The log-likelihood ratio then simplifies to: 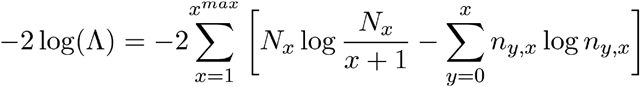. Finally, we use the theorem that the log-likelihood ratio follows a chi-square distribution 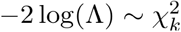, with k (the number of degrees of freedom) determined as the difference between the number of unknown parameters of the null and alternative parameters^43^. In our case, 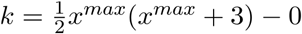. This enables us to perform our likelihood ratio test. We calculated the log-likelihood ratio in Python and utilized the scipy.stats.chi2.sf function to determine the p-value of our test statistic.

## Supplemental figure legends

**Supplemental Figure 1.**
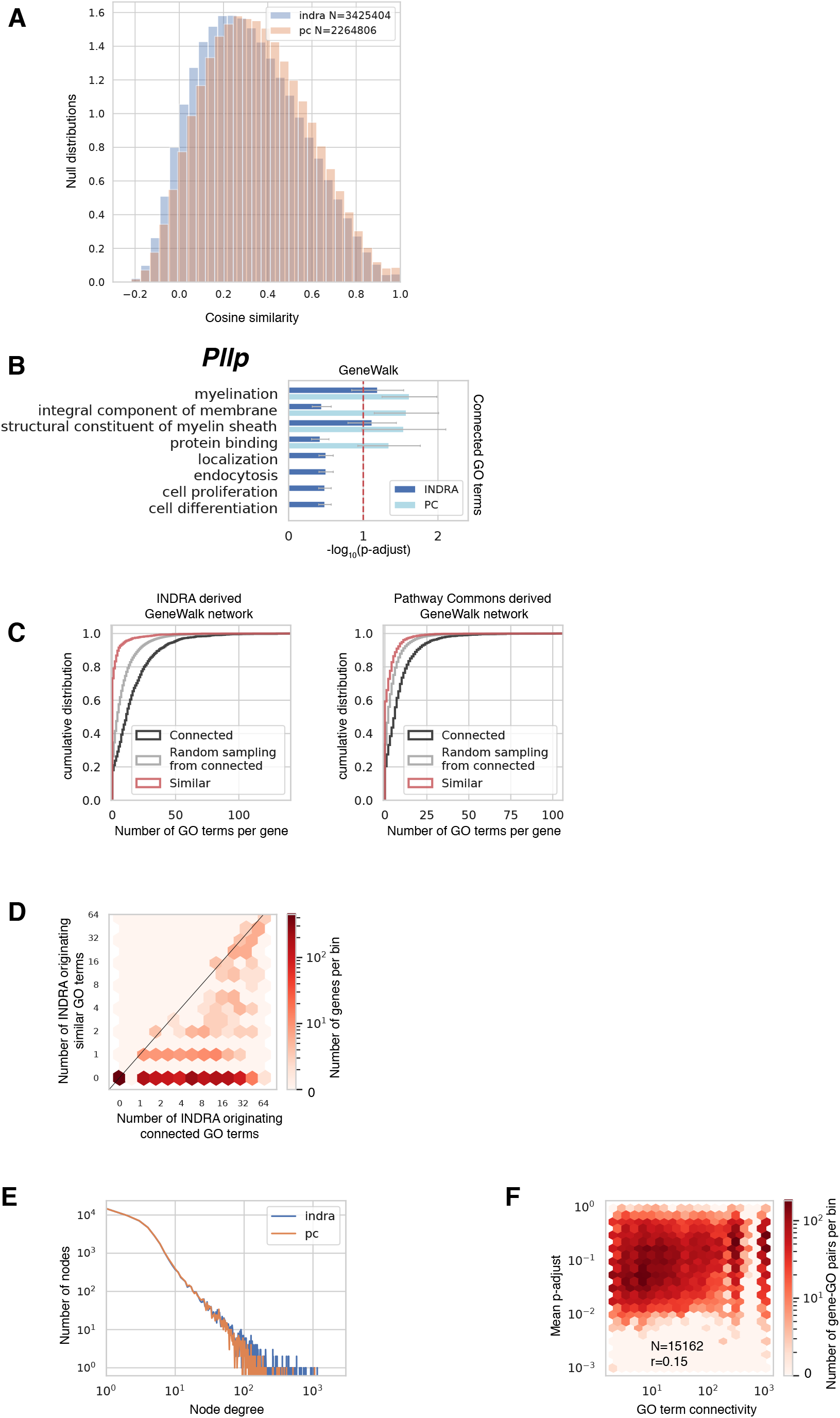
**A** Histograms of cosine similarity value null distributions from the GeneWalk analyses in the *qki* condition with INDRA and Pathway Commons (PC) as knowledge bases. **B** GeneWalk results for *Pllp* in the *Qki*-deficient condition using either INDRA or PC as a knowledge base source to assemble the GeneWalk network (GWN). All GO terms connected to *Pllp* are rank-ordered by Benjamini-Hochberg FDR-adjusted p-value (p-adjust), indicating their functional relevance to *Mal* in the context of *Qki* deletion in oligodendrocytes. Error bars indicate 95% confidence intervals of mean p-adjust. FDR=0.1 (dashed red line) is also shown. Supplemental Tables 1 (INDRA) and 2 (PC) show full GeneWalk results. **C** Cumulative distribution of number of connected (black) and similar (red) GO terms per gene, alongside a simulation that uniformly randomly sampled from the number of connected terms (grey) for GWNs with INDRA (left) and PC (right). **D**. Hexagon density plot for all DE genes (N=1861) in terms of number of INDRA-originating connected GO terms and number of INDRA-originating similar GO terms (at FDR=0.05). **E** Node degree (number of neighboring nodes) distribution function of *qki* GWNs with INDRA (blue) or PC (orange). **F** Hexagon density plot of all tested gene-GO pairs (N= 15162) as a function of GO term connectivity and similarity significance (p-adjust) resulting from the *Qki*-condition GeneWalk using Pathway Commons as a knowledge base. Pearson correlation coefficient r=0.15 is also shown.

**Supplemental Figure 2.**
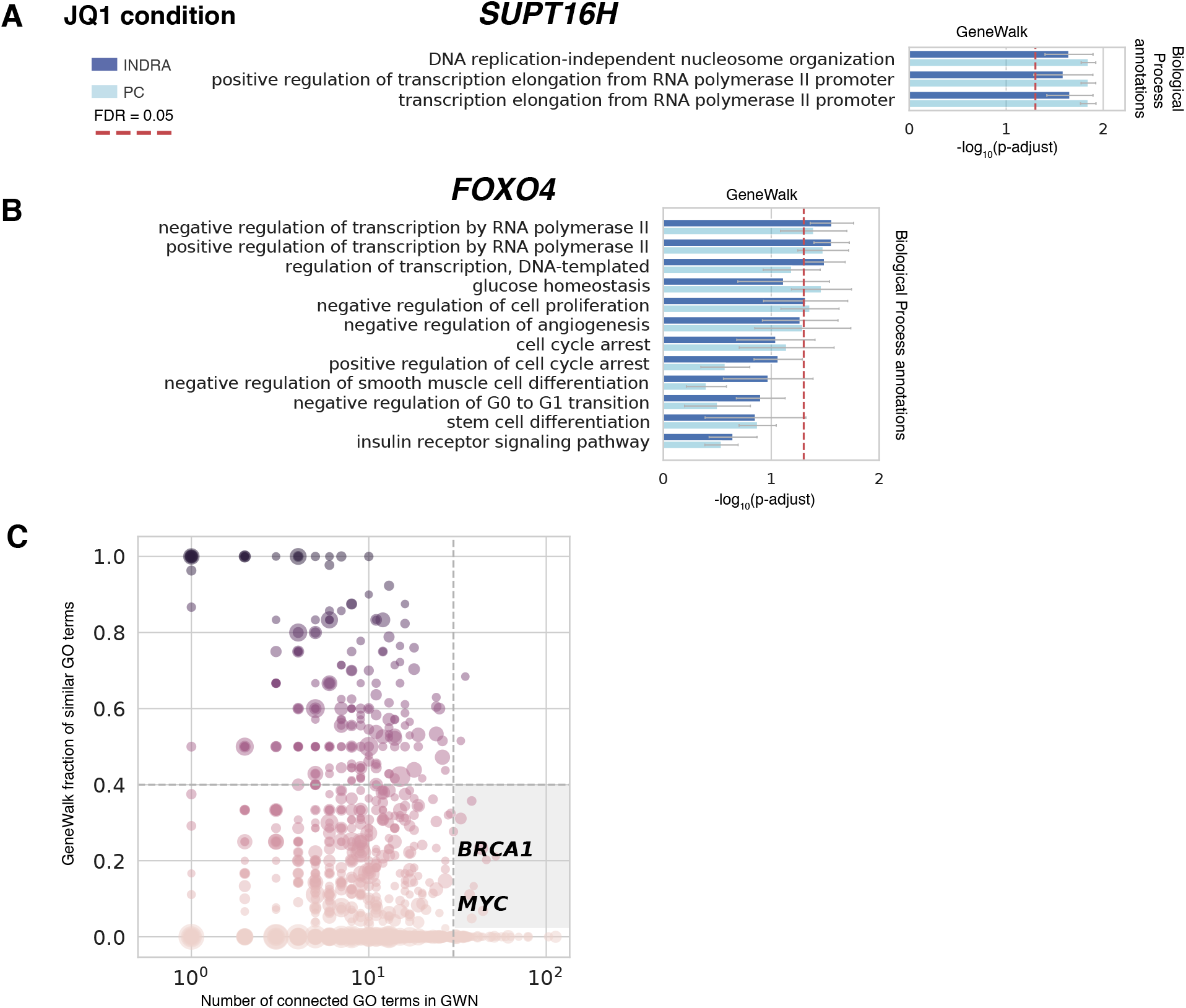
**A** GeneWalk results for the transcriptional regulator *SUPT16H* under JQ1 treatment. Annotated biological processes are rank-ordered by mean FDR adjusted p-value. Error bars indicate 95% confidence intervals of mean p-adjust. FDR=0.05 (dashed red line) is also shown. See Supplemental Table 3 (INDRA) and 4 (PC) for full details. **B** As in (**A**) for *FOXO4*. **C** Scatter plot with DE genes as data points showing the fraction of similar GO terms over total number of connected GO terms (min_f, minimum value between INDRA and PC GWNs) as a function of its number of GO connections (N^GO^). The circle size scales with the differential expression significance strength (-log_10_(p-adjust)) and the color hue with min_f. Nine genes were identified with min_f < 0.4 and N^GO^ > 30 (see Supplemental Table 7 for complete gene list).

**Supplemental Figure 3.**
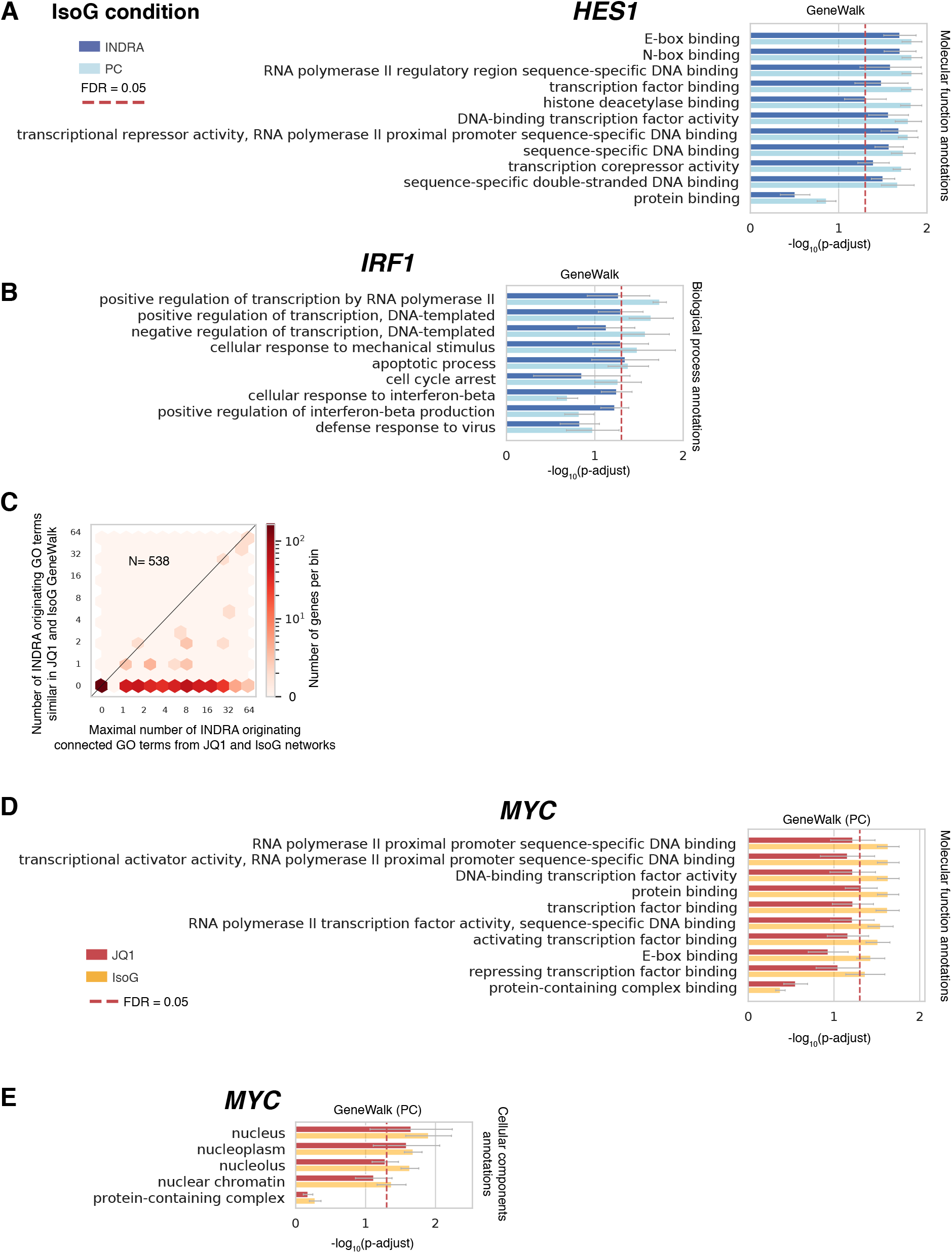
**A** GeneWalk results with INDRA or Pathway Commons (PC) knowledge base for DE gene *HES1*. Annotated molecular function terms are rank-ordered by FDR-adjusted p-value to their indicate relative importance in the IsoG condition. FDR=0.05 (dashed red line) is also shown. See Supplemental Table 5 (INDRA) and 6 (PC) for full details. **B** As in (**A**) for *IRF1* but for biological process terms. **C** Hexagon density plot for overlapping DE genes (N=538) in terms of number of overlapping INDRA originating similar GO terms and number of INDRA originating possible shared connected GO terms. **D-E** GeneWalk results (with PC as data source) for *MYC* in the JQ1 (red) and IsoG (yellow) condition. Annotated molecular function (**D**) or cellular components (**E**) annotations are rank-ordered by FDR-adjusted p-value to their indicate relative importance under the IsoG condition. See Supplemental Table 4 (JQ1) and 6 (IsoG) for full details. Red dashed line indicates FDR=0.05.

## Supplemental Table legends

Supplemental Table 1: GeneWalk output using INDRA as the knowledge base for differentially expressed genes from the *Qki* study.

Supplemental Table 2: GeneWalk output using Pathway Commons as the knowledge base for differentially expressed genes from the *Qki* study.

Supplemental Table 3: GeneWalk output using INDRA as the knowledge base for differentially expressed genes from the JQ1 study.

Supplemental Table 4: GeneWalk output using Pathway Commons as the knowledge base for differentially expressed genes from the JQ1 study.

Supplemental Table 5: GeneWalk output using INDRA as the knowledge base for differentially expressed genes from the IsoG study.

Supplemental Table 6: GeneWalk output using Pathway Commons as the knowledge base for differentially expressed genes from the IsoG study.

Supplemental Table 7: Nine genes identified in JQ1 condition with more than 30 GO annotations of which at most 40% were similar.

## Acknowledgements

This work was supported by National Institutes of Health grant 5R01HG007173-07 (L.S.C.), EMBO fellowship ALTF 2016-422 (R.I.), and DARPA grants W911NF-15-1-0544 and W911NF018-1-0124 (P.K.S.). We thank Karine Choquet and Heather Drexler for advice on the previously described mouse RNA-seq differential expression^23^ analysis and IsoG NET-seq^36^. We thank Dylan Marshall and other members of the Churchman lab for discussions.

## Author contributions

R.I. and L.S.C. conceived the study. R.I. developed the GeneWalk methodology. R.I. and B.G. implemented and released the GeneWalk software. B.G., J.B. and P.K.S. developed INDRA and provided the reaction statements. R.I. performed data analysis. R.I. and L.S.C wrote the manuscript with input from all authors.

